# Microfluidic Mechanical Reactivation of Aged Stem Cells

**DOI:** 10.64898/2026.02.25.707893

**Authors:** Soo Bin Jang, Tak-Il Jeon, Geun-Ho Kang, Dongwhan Seo, Hyelee Kim, Hancheol Yeo, Jaekwon Seok, Kyung Min Lim, Ahmed Abdal Dayem, Se Jong Kim, Kwonwoo Song, Yeonjoo Kwak, Jeongsoo Hur, Aram J. Chung, Ssang-Goo Cho

**Affiliations:** Department of Stem Cell and Regenerative Biotechnology, School of Advanced Biotechnology, Molecular & Cellular Reprogramming Center, Institute of Advanced Regenerative Science, and Institute of Health, Aging & Society, Konkuk University, Seoul 05029, Republic of Korea; R&D Team, StemExOne CO., Ltd., 19, Achasan-ro 5-gil, Seongdong-gu, Seoul 04793, Republic of Korea; School of Biomedical Engineering and Interdisciplinary Program in Precision Public Health, Korea University, Seoul 02841, Republic of Korea; MxT Biotech, Seoul 04785, Republic of Korea

**Keywords:** Microfluidics, Mechanobiology, Stem cell reactivation, Cytoskeletal remodeling, Partial rejuvenation

## Abstract

Stem cell aging significantly impairs therapeutic efficacy, requiring innovative strategies to restore potency. We present a microfluidic cell-compressing platform for reactivation (μ-CPR) designed to apply controlled hydrodynamic deformation to late-passage stem cells. This mechanical stimulation facilitates functional reactivation without external chemical cues. Within a defined window, μ-CPR effectively reduces oxidative stress, SA-β-Gal activity, and γH2AX foci, while simultaneously restoring proliferation and canonical stemness markers (OCT4, SOX2, and KLF4). Mechanical stimulation via μ-CPR induces coordinated structural remodeling: nuclei become more compact, actin cortex organization is restored, α-actinin redistributes to focal adhesions, and microtubule networks are restructured, suggesting a rebalanced intracellular tension. Transcriptomic and proteomic analyses reveal that this process reprograms extracellular matrix remodeling and DNA repair pathways while attenuating pro-fibrotic and senescence-associated secretory phenotype (SASP)-associated pathways. Crucially, this reactivation occurs without compromising fundamental MSC hallmarks, preserving intrinsic immunophenotypes and multilineage differentiation potential. Functionally, μ-CPR-processed stem cells demonstrate restored *in vitro* wound closure and enhanced tissue repair *in vivo*, with efficacy appearing dependent on mechanical dosage. This platform establishes a non-genetic, mechanobiological approach to restoring stem cell function, offering a scalable strategy for functional reactivation and potentially paving the way toward comprehensive cellular rejuvenation.

## 1. Introduction

Cells are intrinsically mechanosensitive and continuously integrate biophysical cues from their microenvironment to regulate their fate and function.^1^ Mechanical forces, including shear stress, matrix elasticity, cyclic strain, and compressive loading, govern proliferation, differentiation, migration, and apoptosis.^2–6^ Through mechanotransduction, which involves transforming mechanical inputs into biochemical signals, cells coordinate tissue morphogenesis, homeostasis, and regeneration, establishing the foundation of mechanobiology.^4, 7, 8^ These insights have motivated the development of various experimental systems, including deformable substrates, micropillar arrays, atomic force microscopy, and microfluidic devices, to probe the molecular pathways underlying cellular mechanotransduction.^7, 9^ Collectively, these studies have uncovered pathways through which cells sense and respond to mechanical stimuli, providing a quantitative framework to understand the force-response relationship at the molecular and cellular levels.^4, 9, 10^

Building on these findings, mechanical stimulation has evolved from an observed biophysical phenomenon into a precise engineering tool for the strategic manipulation of cellular behavior.^11^ Externally applied forces can guide stem cell lineage commitment^12^, enable non-viral intracellular cargo delivery^13^, and regulate immune cell activation.^14^ These approaches highlight the potential of mechanical inputs as non-chemical regulators of cell state, offering distinct advantages over conventional biochemical methods.^15^ However, while most applications have focused on guiding differentiation, activation, or transient priming, the feasibility of utilizing mechanical stimulation to reactivate aged stem cells by specifically restoring youthful phenotypes and mitigating senescence-associated dysfunction, remains largely unexplored.^16^

This gap in knowledge is particularly critical because stem cell aging is characterized by diminished self-renewal, loss of multipotency, accumulation of reactive oxygen species (ROS), epigenetic drift, and the acquisition of senescence-associated secretory phenotype (SASP).^17, 18^ *In vitro* expansion accelerates these effects, where conditions such as supraphysiological oxygen tension and the loss of niche-derived cues induce replicative stress, oxidative damage, and chromatin remodeling.^19–21^ These changes manifest as nuclear enlargement, cytoskeletal stiffening, and reduced mechanosensory plasticity, factors that restrict regenerative ability and compromise the scalability of stem cell-based therapeutics.^18, 22^ In this context, developing effective strategies that can restore the early-passage transcriptional, epigenetic, and functional states of aged or senescent cells, even if only partially or transiently, would represent a significant advancement for clinical translation.^18, 23^ In alignment with this potential, recent reports suggest that mechanical cues can effectively attenuate senescence markers and at least transiently restore the proliferative and functional capacity of stressed or aged cells.^16, 24^ However, the broader application of these findings is currently hindered by the limitations of existing systems, which primarily utilize indirect modulation or low-throughput modalities, such as substrate stiffness tuning or cyclic stretching.^25^ These conventional approaches often lack the mechanistic specificity, reproducibility, and scalability essential for clinical translation.^16, 26^ Consequently, a pressing need remains for advanced mechanostimulation platforms capable of delivering controlled, high-throughput, and physiologically relevant forces to facilitate the functional reactivation of aged stem cells without the inherent complexities of genetic or pharmacological intervention.^16, 26^

To address these challenges, we developed a high-throughput microfluidic cell-compressing platform for reactivation (μ-CPR) (**Figure 1A**), which subjects cells to uniform hydrodynamic deformation (**Figure 1B**). By applying a defined mechanical strain within a precise temporal window, this platform facilitates the restoration of cellular functions without chemical intervention. Our findings demonstrate an attenuation of senescence-associated β-galactosidase (SA-β-Gal) activity, diminished ROS levels, and the restoration of proliferation and multipotency markers, alongside a transcriptomic reversion toward early-passage states across multiple stem cell types. Functionally, μ-CPR-treated cells exhibit enhanced regenerative performance in *in vitro* scratch assays and *in vivo* wound-healing models (**Figure 1C**). Furthermore, mechanistic analyses reveal a coordinated activation of DNA repair and redox-balancing pathways concurrent with the inhibition of pro-inflammatory networks, effectively linking cytoskeletal-nuclear remodeling to functional reactivation. Collectively, μ-CPR presents a scalable, reproducible, and mechanistically distinct approach for restoring stem cell potency, with broad implications for next-generation regenerative medicine.

**Figure 1.**
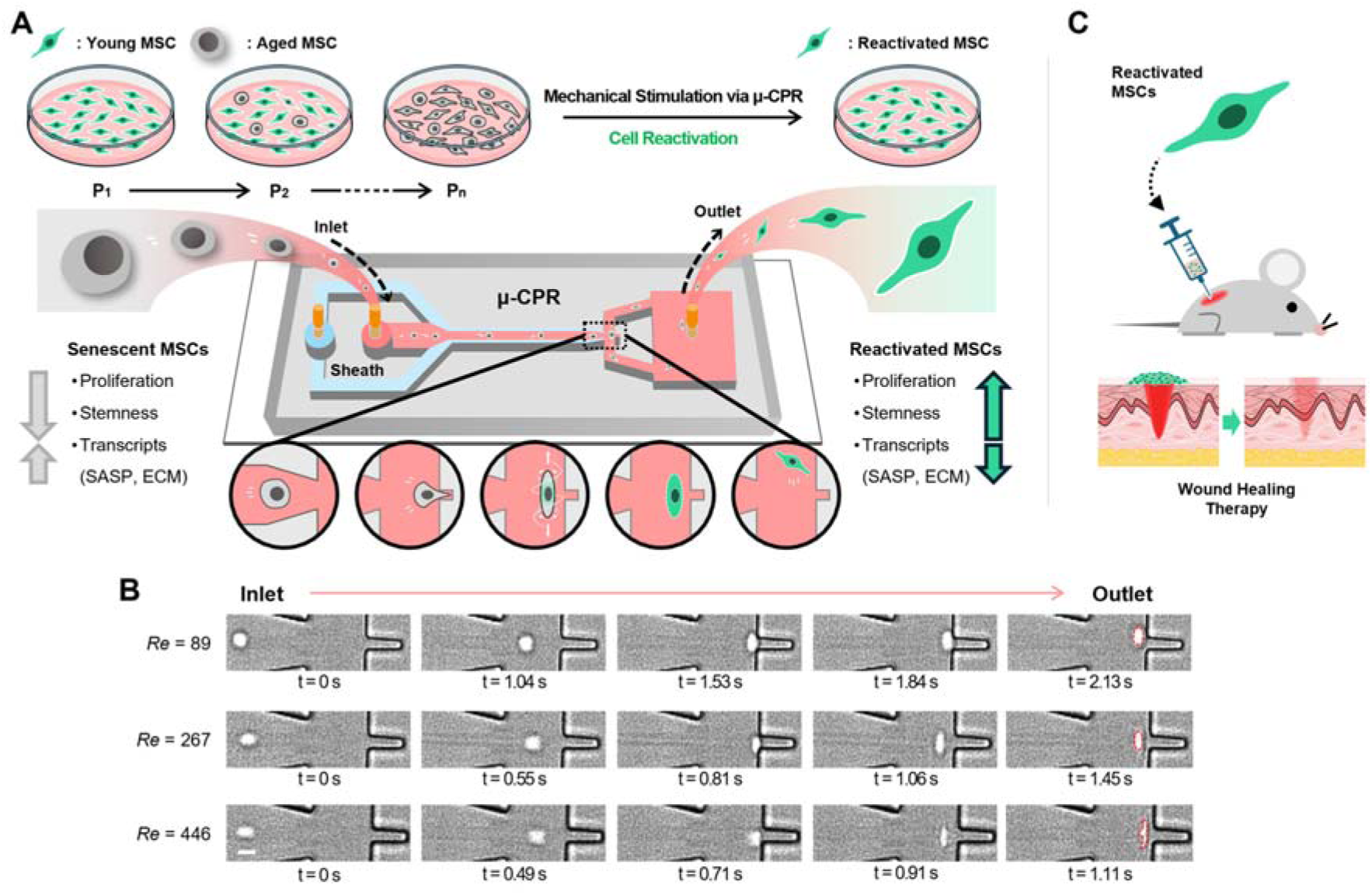
Mechanical stimulation via the μ-CPR platform to functionally reactivate aged mesenchymal stem cells (MSCs) and enhance wound healing. **(A)** Schematic of MSC reactivation using the μ-CPR platform. **(B)** High-speed microscopy images showing flow-rate-dependent changes in cell morphology during hydrodynamic cell compression. Scale bar = 10 μm. **(C)** Schematic illustrating the wound healing progression facilitated by μ-CPR-reactivated MSCs.

## 2. Results and Discussion

### 2.1 Morphological and Proliferative Responses to μ-CPR

The μ-CPR device builds on our previous microfluidic platform, designed for the intracellular delivery of exogenous nanomaterials, such as nucleic acids and nanoparticles, into immune cells. While the overall architecture appears similar, the channel geometry and fluidic conditions were strategically re-engineered to accommodate the larger size and distinct mechanical properties of stem cells. These modifications enable controlled stem cell deformation while maintaining high cell viability, providing a tunable platform for delivering physiologically relevant mechanical cues.

Given its ability to apply uniform mechanical stress to single cells at high throughput, we hypothesized that μ-CPR could function as a mechanical reactivation tool for aged stem cells. As a representative model, we first examined Wharton’s Jelly-derived MSCs (WJ-MSCs), known for their immunomodulatory properties and potential in treating immune-mediated disorders. Moreover, their postnatal origin circumvents the ethical limitations associated with embryonic stem cell sources, making them ideal for translational research.

Fluidically induced deformation was evaluated by subjecting WJ-MSCs to μ-CPR under varying flow rates, characterized by the Reynolds number (*Re*; a dimensionless number describing the ratio between inertial and viscous forces). As expected, cell elongation increases proportionally with *Re* (**Figure 1B**). We then evaluated whether this controlled mechanomodulation influences proliferative capacity, given that aged MSCs typically exhibit slower proliferation than early-passage cells. Late-passage WJ-MSCs (P18) exhibit significantly enhanced proliferation following μ-CPR treatment compared to unstimulated controls (**Figure 2A**). Notably, proliferation declined sharply when *Re* exceeded 267, indicating a threshold beyond which excessive deformation compromises membrane integrity and recovery. This phenomenon likely reflects transient membrane permeabilization and delayed resealing at higher mechanical loads, resulting in reduced viability.

**Figure 2.**
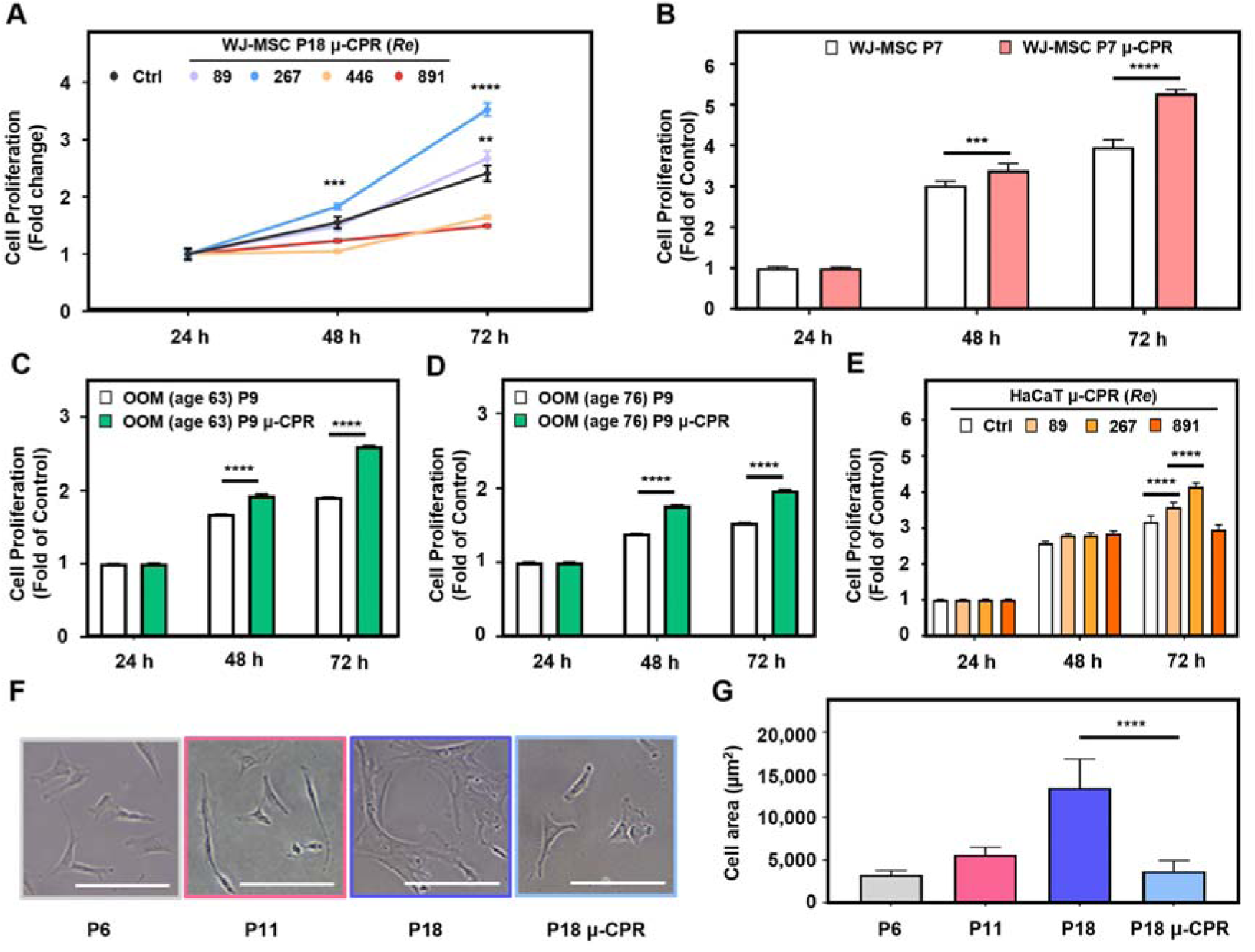
μ-CPR enhances proliferative activity and attenuates senescence-associated morphological changes in MSCs. **(A)** Time-dependent proliferation analysis of senescent Wharton’s Jelly-derived MSCs (WJ-MSCs) stimulated under various μ-CPR conditions (*Re* = 89, 267, 446, and 891). **(B)** Comparative analysis of WJ-MSCs (P7) proliferation with and without μ-CPR stimulation (*Re* = 267). **(C, D)** Proliferation analysis of orbicularis oculi muscle-derived stem cells (OOM-SCs) isolated from donors aged 63 and 76 years with and without μ-CPR (*Re* = 267). **(E)** Proliferation analysis of immortalized human keratinocytes (HaCaT cells) under various μ-CPR conditions (*Re* = 89, 446, and 891). **(F)** Cell-size comparison among early- (P6), mid- (P11), and late-passage (P18) WJ-MSCs, alongside μ-CPR-treated senescent WJ-MSCs (P18 μ-CPR). Scale bar = 200 µm. **(G)** Quantification of cell size using ImageJ. Data represent the mean ± SEM of three independent experiments. Statistical analysis was performed using one-way ANOVA; \**P* < 0.05, \*\**P* < 0.01, \*\*\**P* < 0.001, \*\*\*\**P* < 0.0001.

Encouraged by the enhanced proliferation, we assessed whether this effect extends across different cellular states and origins. Early-passage WJ-MSCs (P7) also demonstrate increased proliferation after μ-CPR (**Figure 2B**), confirming that the mechanical activation consistently augments proliferative activity across passage numbers. Similarly, aged human orbicularis oculi muscle (OOM)-derived MSCs display enhanced proliferation following μ-CPR stimulation (**Figures 2C–D**), indicating that the effect is not limited to a single tissue source. To validate these findings, we tested immortalized human keratinocytes (HaCaT) and observed a similar increase in proliferation post-treatment (**Figure 2E**). These results collectively indicate that μ-CPR-induced mechanomodulation promotes proliferative activation across diverse cell types and ages.

In addition, we analyzed morphological features associated with cellular senescence. Aged WJ-MSCs exhibit enlarged, flattened morphologies consistent with cytoskeletal stiffening. Following μ-CPR treatment, we observed a significant reduction in cell size, as shown in **Figure 2F**. Moreover, quantitative analysis at 72 h reveals a significant decrease in mean cell area (**Figure 2G**). Furthermore, time-resolved measurements from 0.5 to 8 h post-treatment (**Figure S1**) confirm dynamic actin reorganization suggestive of a shift toward a more youthful state. Because cellular enlargement is a characteristic feature of senescence, and prior studies have shown that mechanical stretching can restore the proliferative capacity in aged MSCs and keratinocytes, our findings suggest that μ-CPR-treated cells demonstrate the phenotypic and functional properties of cellular reactivation.

### 2.2 Molecular and Molecular Analysis of μ-CPR–Treated Stem Cells

Having established that μ-CPR induces proliferative and morphological modulations at the macroscopic cellular level, we investigated whether these effects are accompanied by molecular and structural reprogramming. Cellular aging is characterized by increased SA-β-Gal activity, elevated ROS levels, enhanced γH2AX (the phosphorylated form of H2AX histone family member X) expression, loss of stemness, and altered cellular structures. To determine whether μ-CPR attenuates these hallmarks, we systematically analyzed biochemical and molecular markers associated with senescence in late-passage stem cells.

As size reduction was confirmed previously, we first assessed ROS levels in aged WJ-MSCs (P18). Aged MSCs typically exhibit elevated ROS due to impaired mitochondrial function and weakened antioxidant defenses. The DCF-DA fluorescence assay revealed that μ-CPR treatment significantly decreased ROS levels (**Figure 3A**) in a flow rate-dependent manner (**Figure 3B**). Moreover, moderate mechanical stimulation significantly reduced intracellular ROS, whereas excessive deformation (*Re* >267) diminished cell viability, consistent with the threshold-dependent effects observed in proliferation assays. These findings indicate that controlled hydrodynamic deformation mitigates oxidative stress within an optimal mechanical window.

**Figure 3.**
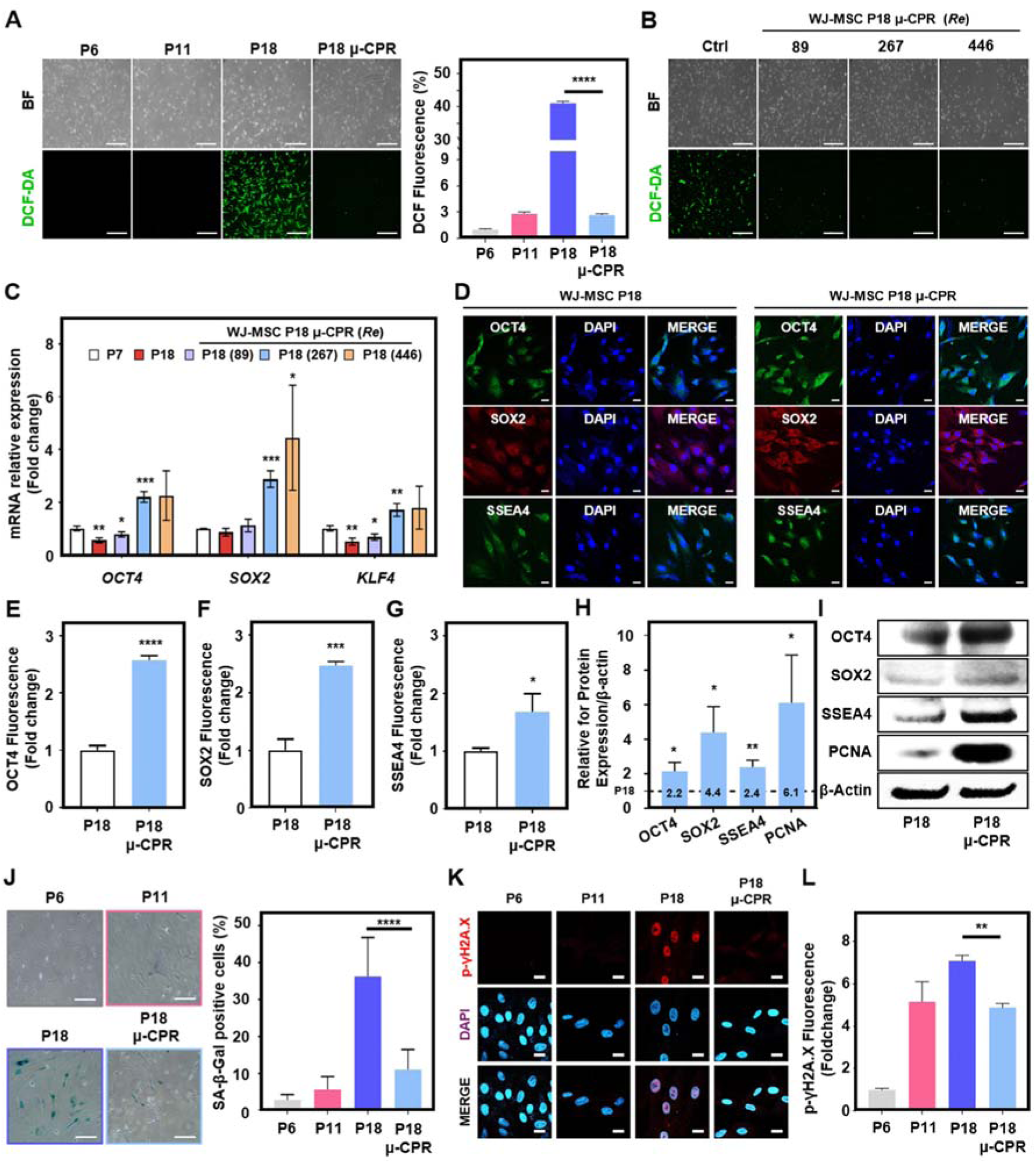
μ-CPR reduces reactive oxygen species (ROS), attenuates senescence markers, and restores stemness in late-passage MSCs. **(A)** Representative fluorescence images and comparative analysis of intracellular ROS levels among P6, P11, P18, and P18 μ-CPR cells. Scale bar = 500 µm. **(B)** Representative fluorescence images showing intracellular ROS levels in P18 μ-CPR under three flow conditions (*Re* = 89, 267, and 446). Scale bar = 500 µm. **(C)** RT-qPCR analysis of stemness-related genes (*OCT4*, *SOX2*, and *KLF4*) after 3 days. Expression levels are normalized to P7 WJ-MSCs and presented as relative ratios for the P18 μ-CPR groups (*Re* = 89, 267, and 446). **(D)** Representative immunofluorescence images showing the expression of stemness markers (OCT4, SOX2, and SSEA4) in P18 and P18 μ-CPR cells. Nuclei were stained with DAPI. Scale bar = 20 µm. **(E–G)** Quantification of fluorescence intensity (fold change) for OCT4 **(E),** SOX2 **(F)**, and SSEA4 **(G)** in P18 and P18 μ-CPR cells. **(H)** Quantification of Western blot band intensities (fold change) for stemness markers (OCT4, SOX2, and SSEA4) and the proliferation marker PCNA in P18 μ-CPR cells relative to the P18 control. **(I)** Immunoblotting of OCT4, SOX2, SSEA4, PCNA, and β-actin in μ-CPR and control MSCs. **(J)** Relative quantification of senescent cells using SA-β-Gal staining in P6, P11, P18, and P18 μ-CPR cells. The graph presents the percentages of SA-β-Gal-positive cells. Scale bar = 500 µm. **(K, L)** Evaluation of intracellular γH2AX expression and its quantification in P6, P11, P18, and P18 μ-CPR cells. Scale bar = 20 µm. Data are presented as the mean ± SEM of three independent experiments. Statistical analysis was performed using one-way ANOVA (\**P* < 0.05, \*\**P* < 0.01, \*\*\**P* < 0.001, \*\*\*\**P* < 0.0001).

Next, we examined whether μ-CPR restores stem cell identity by quantifying the expression of canonical pluripotency-associated transcription factors, namely OCT4, SOX2, and KLF4, via RT-qPCR and immunofluorescence. As expected, early-passage WJ-MSCs exhibited higher expression of these markers compared with late-passage cells (**Figure 3C**). Following μ-CPR treatment, transcript levels in aged WJ-MSCs were significantly upregulated to approximately two to four times those of early-passage controls. Immunofluorescence imaging confirmed enhanced nuclear localization of OCT4, SOX2, and KLF4 (**Figure 3D**), with fluorescence intensities increasing by 1.8- to 2.5-fold in μ-CPR-treated cells (**Figures 3E–H**). Immunoblotting validated these findings, revealing increased expression of OCT4, SOX2, SSEA4, KLF4, and the proliferation marker, proliferating cell nuclear antigen (PCNA) (**Figure 3I**). Collectively, these results indicate the reactivation of stemness-associated transcriptional programs.

To evaluate senescence attenuation directly, we measured SA-β-Gal activity. Late-passage WJ-MSCs showed a pronounced reduction in the number of SA-β-Gal-positive cells after μ-CPR treatment (**Figure 3J**). Quantitative analysis revealed that the proportion of senescent cells decreased to levels comparable with early-passage controls (P11), demonstrating more than a twofold reduction relative to untreated P18 cells. Similar declines in SA-β-Gal activity were observed across diverse aged cell types, including umbilical cord-derived MSCs, adipose-derived stem cells, OOM-derived stem cells, and human fibroblasts (BJ cells), which are non-stem cells. In all cell types tested, μ-CPR treatment consistently reduced SA-β-Gal activity (**Figure S2**), demonstrating its generalizability.

We further investigated DNA damage response signaling by analyzing γH2AX, a marker of DNA double-strand breaks. As expected, γH2AX levels were elevated in aged WJ-MSCs, reflecting accumulated genotoxic stress. Following μ-CPR treatment, γH2AX nuclear foci substantially decreased, as shown by immunofluorescence (**Figure 3K**), with quantification confirming significantly reduced fluorescence intensity (**Figure 3L**). These results suggest that controlled mechanical stimulation suppresses DNA damage-associated signaling, thereby attenuating senescence-associated pathways.

Because senescence is linked to altered nuclear architecture and cytoskeletal organization, we next assessed the structural remodeling that occurs after μ-CPR treatment. Quantitative image analysis revealed that nuclei of late-passage WJ-MSCs were enlarged and irregular, indicative of chromatin relaxation and loss of nuclear tension,^27, 28^ whereas μ-CPR treatment restored compact nuclear morphology comparable to that of early-passage cells. In P18 cells, μ-CPR significantly reduced the nuclear major axis at 48 h (*P* < 0.01) and 72 h (*P* < 0.001) relative to untreated late-passage cells (**Figure S3A**). Concomitantly, senescent MSCs exhibited disrupted cytoskeletal organization: actin filaments formed thick, disordered stress fibers, α-actinin was delocalized from focal adhesions, and microtubule networks were fragmented (**Figure S3B–D**). In contrast, μ-CPR-treated cells displayed a well-organized actin cortex, restored α-actinin networks at focal adhesions, and restructured microtubule networks, indicating that μ-CPR reestablishes cytoskeletal-nuclear connectivity, facilitating effective force transmission and maintaining cellular polarity, which are crucial for stem cell function and regenerative capacity.^29, 30^

Based on these reactivation characteristics, we next aimed to verify whether the μ-CPR platform affects the innate traits of MSCs. First, we measured the levels of MSC-specific surface markers. Flow cytometric analysis revealed that the expression levels of these surface markers remained unaltered following μ-CPR treatment, confirming that the mechanical stimulation stably preserves the intrinsic immunophenotype of the stem cells (**Figure S4A**). Beyond surface marker analysis, we then investigated multilineage differentiation potential after μ-CPR treatment. As shown in **Figure S4B**, μ-CPR-treated cells maintained their differentiation potential toward adipogenic and osteogenic lineages at levels comparable to those of early-passage controls. To further validate this, the colony-forming unit-fibroblast (CFU-F) efficiency, a key indicator of self-renewal capacity, was characterized. The μ-CPR treated group exhibited a more than twofold increase compared to the aged control group (**Figure S4C**). These results indicate that μ-CPR enhances functional capacity without compromising the defining phenotypic identity of MSCs.

To further corroborate these observations at the molecular level, we performed RT-qPCR analysis of major senescence-associated markers (**Figure S4D**). The expression levels of *p16* (*CDKN2A*), *p21* (*CDKN1A*), and *p53* (*TP53*), which are characteristically upregulated in senescent cells, were significantly downregulated following μ-CPR treatment. Notably, these qPCR results were highly consistent with our transcriptomic analysis (discussed in detail in the following section), which also demonstrated a marked reduction in these senescence-associated genes (**Figure S5**). This validation underscores the robustness of μ-CPR in suppressing the molecular program of cellular senescence.

These findings collectively demonstrate that μ-CPR partially attenuates multiple traits of cellular aging, including oxidative stress, senescence marker expression, DNA damage signaling, and cytoskeletal disorganization, while restoring stemness and proliferative potential. Thus, μ-CPR serves as a mechanistically distinct, non-genetic, and broadly applicable strategy for functional reactivation of aged stem cells without altering their fundamental identity.

### 2.3 Transcriptomic Analysis of μ-CPR–Treated Stem Cells

To elucidate how transient mechanical stimulation via μ-CPR reprograms senescent transcriptional networks, we performed transcriptomic profiling using RNA sequencing (RNA-seq). RNA-seq analysis of μ-CPR-treated late-passage WJ-MSCs revealed global expression patterns indicative of partial functional restoration. Compared with untreated late-passage cells, μ-CPR-treated cells exhibited a substantially smaller set of differentially expressed genes (DEGs), consisting of 26 upregulated and 115 downregulated transcripts (**Figure 4A**). Among the upregulated genes, *MMP1*, *CSF3*, *PLAU*, *MMP3*, *IL33*, *SERPINB2*, and *THBD* were enriched in pathways related to extracellular matrix (ECM) degradation, fibrinolysis, and cytokine signaling, whereas downregulated transcripts primarily mapped to collagen biosynthesis and structural ECM organization (**Figures 4B–C**). These findings suggest that μ-CPR suppresses pro-fibrotic and SASP-associated programs^31, 32^ while enhancing tissue remodeling capacity, thereby mitigating the inflammatory and structural constraints that perpetuate senescence. Because the ECM and cytokine milieu directly regulate nuclear mechanotransduction and chromatin accessibility, these changes likely contribute to transcriptional reprogramming toward a more youthful phenotype.

**Figure 4.**
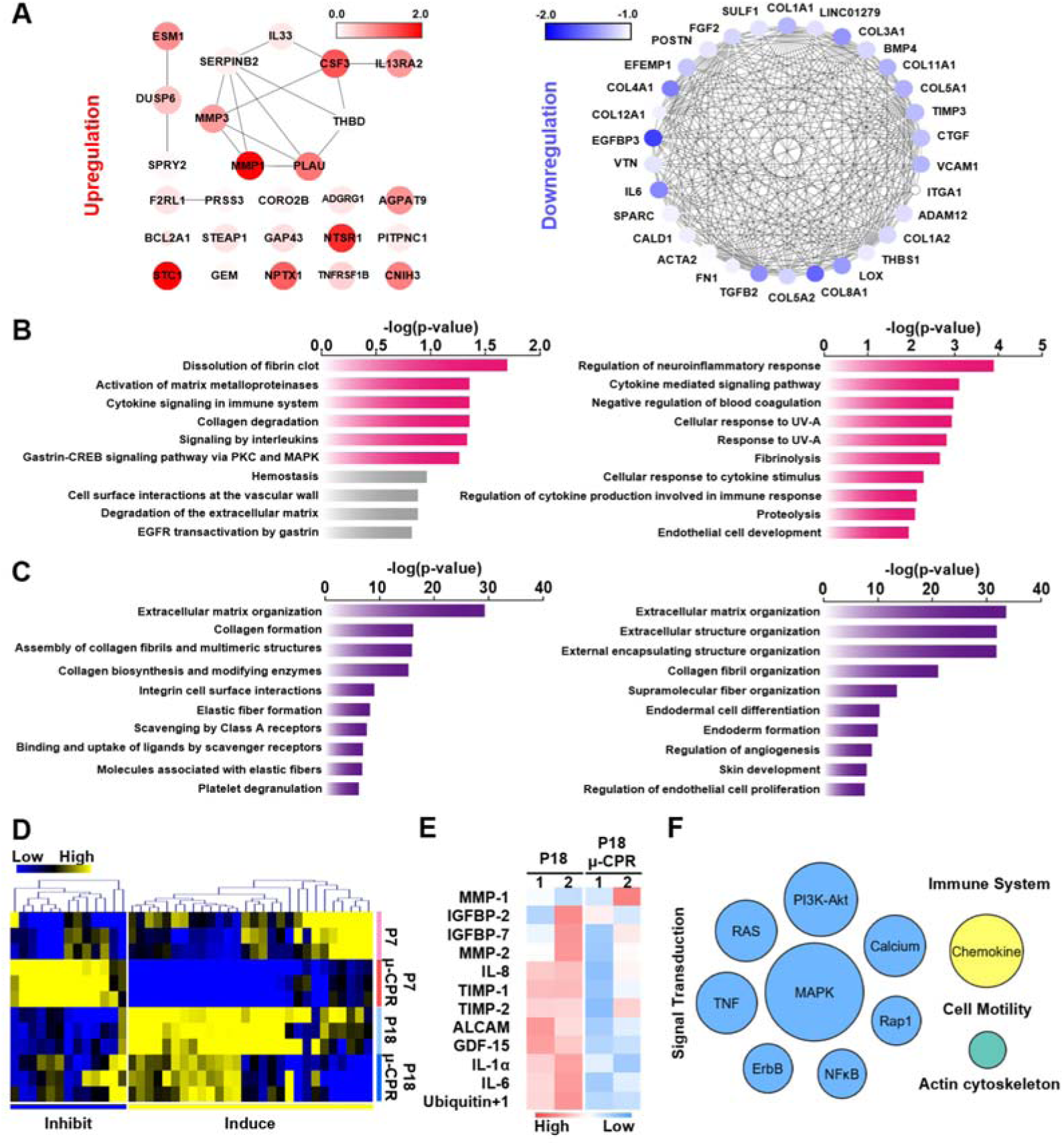
Bioinformatic analysis of senescent WJ-MSCs subjected to. μ**-CPR. (A)** Protein interaction maps showing differentially expressed genes (DEGs) in senescent WJ-MSCs following μ-CPR treatment compared with control cells. Node saturation indicates the magnitude of gene expression differences, while node size reflects the statistical significance (*P*-value). Bar graphs showing predicted pathways enriched in **(B)** upregulated and **(C)** downregulated genes, ranked by *P*-value based on Gene Ontology and Reactome pathway analyses. **(D)** Hierarchical clustering heatmaps showing expression trends of senescence-associated and senescence-inhibited genes in P7 and P18 cells relative to controls, with colors indicating gene expression differences according to z-score thresholds. **(E)** Hierarchical clustering heatmaps showing expression patterns of key SASP-related proteins in P18 and P18 μ-CPR groups, with colors indicating protein expression differences according to z-score thresholds. **(F)** The top 20 cytokines identified from the protein array were enriched in specific signaling pathways determined using the KEGG database.

Heatmap analyses of aging-related gene modules (**Figures 4D** and **S5**) reinforced this trend. Expression signatures associated with senescence and aging were broadly diminished, whereas those linked to longevity and genome maintenance, such as base excision repair, nucleotide excision repair, and double-strand break repair, were upregulated. This shift aligns with the reduced ROS accumulation and γH2AX foci observed experimentally,^33, 34^ highlighting the correspondence between transcriptomic and phenotypic indicators of improved DNA repair capacity.^35, 36^ Furthermore, μ-CPR elevates the expression of DNA repair and senescence-suppressive genes while repressing canonical senescence markers, including *CDKN2A*, *CDKN1A*, and *IL6*. These transcriptional shifts coincided with a decrease in SA-β-Gal-positive cells and the restoration of OCT4, SOX2, and KLF4 expression^37, 38^, indicating enhanced stemness.

Integration with protein-level analyses further corroborated these findings. Cytokine array profiling revealed reduced secretion of IL-6, IL-8, MMP-1, and MMP-2, accompanied by increased levels of TIMP-1 and TIMP-2 in μ-CPR–treated cells (**Figures 4E–F**), indicating effective suppression of SASP^39, 40^ and reinforcement of ECM homeostasis. The transcriptional modulation of ECM-related pathways further complements our imaging data showing nuclear compaction and cytoskeletal reorganization,^41^ which reinforces the mechanotransductive mechanism of μ-CPR. Notably, the limited number of enriched pathways and observed alterations in cell-cycle, p53, and MAPK/PI3K signaling modules (**Figure S4**) indicate that, while μ-CPR restores youthful transcriptional programs, late-passage cells exhibit constrained transcriptomic plasticity compared with early-passage cells. These findings establish μ-CPR as a mechanistically selective intervention that integrates molecular, structural, and functional reprogramming, achieving a coordinated and partial restoration of aging-associated transcriptional programs in late-passage WJ-MSCs.

### 2.4 Enhanced wound-healing effect from μ-CPR-processed WJ-MSCs

MSC-based therapies are extensively studied for wound healing due to their ability to secrete trophic factors, modulate inflammatory responses, and promote tissue regeneration. However, their therapeutic efficacy progressively diminishes with prolonged expansion as cells undergo replicative senescence, limiting clinical application. To overcome this barrier, we hypothesized that mechanically activating senescent MSCs via μ-CPR could restore their functional potency and enhance their wound-healing capacity (**Figure 5A**).

**Figure 5.**
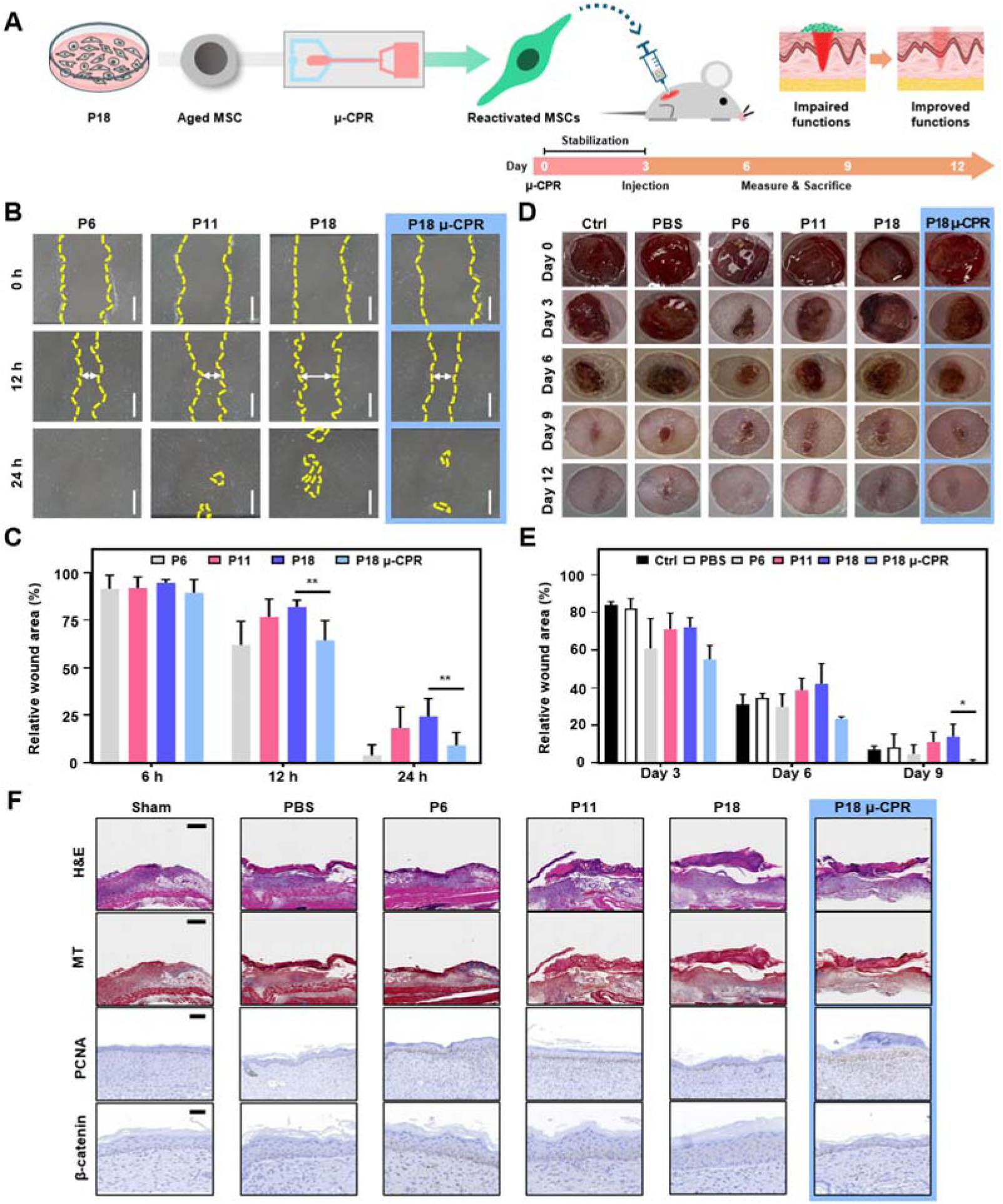
Enhanced *in vivo* wound healing by. μ**-CPR-processed WJ-MSCs. (A)** Schematic representation of *in vivo* wound healing. **(B)** Scratch migration assay comparing P6, P11, and P18 cells. Yellow dotted lines indicate cell-free areas, and white arrows denote migration distance. P18 μ-CPR cells showed increased migratory capacity compared with untreated P18 cells. Scale bar = 500 µm. **(C)** Quantitative results are presented as mean ± SEM from three independent experiments; statistical significance was determined by two-way ANOVA. **(D)** Representative images of wound healing up to day 12 (initial wound diameter: ∼8 mm) for the P6, P11, P18, and P18 μ-CPR groups, and **(E)** quantitative analysis of wound closure at corresponding time points; statistical analyses were performed using the Kruskal-Wallis test. **(F)** Comparative histological analyses of *in vivo* wound healing using P6, P11, P18, and P18 μ-CPR cells, assessed through hematoxylin and eosin (H&E), Masson’s trichrome (MT) staining (day 6, Scale bar = 500 µm), and immunohistochemistry for PCNA (Scale bar = 200 µm) and β-catenin (Scale bar = 20 µm) (day 12). (Control: n = 4; PBS: n = 4; P6: n = 4; P11: n = 4; P18: n = 4; P18 μ-CPR: n = 4; \**P* < 0.05, \*\**P* < 0.01, \*\*\**P* < 0.001, \*\*\*\**P* < 0.0001).

To test this, we compared wound closure rates among early- (P6), mid- (P11), and late-passage (P18) WJ-MSCs, alongside μ-CPR-stimulated senescent cells (P18 μ-CPR). *In vitro* scratch assays demonstrated that μ-CPR treatment substantially enhanced the proliferation and migration of late-passage WJ-MSCs, resulting in wound closure dynamics comparable to early-passage cells (**Figures 5B** and **C**). These findings align with the observed cytoskeletal reorganization and increased proliferative activity following μ-CPR treatment.

Encouraged by these results, we evaluated the regenerative performance of μ-CPR-processed WJ-MSCs *in vivo* using a murine full-thickness skin injury model. Equal numbers of P6, P11, P18, and P18 μ-CPR cells were transplanted into wounds (see Methods for details). At 3 days post-transplant, wound closure rates were similar across groups. However, by day 6, significant differences emerged. By day 9, wound closure in P18 μ-CPR-treated cells matched that of P6-treated cells and exceeded that of untreated P18 cells (**Figures 5D** and **E**), indicating the recovery of therapeutic efficacy following mechanical activation.

To further examine tissue-level repair, wound sections collected at day 6 were analyzed by histological and immunohistochemical staining. Hematoxylin and eosin (H&E) and Masson’s trichrome (MT) staining revealed well-organized neo-epidermis and abundant granulation tissue in the P6 and P18 μ-CPR groups, whereas untreated P18 cells showed limited granulation (**Figure 5F**). PCNA immunostaining indicated a high density of proliferating cells in the P6, P11, and P18 μ-CPR groups, consistent with enhanced cellular proliferation.

Furthermore, β-catenin expression, which declines with cellular aging, was restored in P18 μ-CPR-treated cells, suggesting the reactivation of regenerative signaling pathways (**Figure 5F**). In summary, μ-CPR treatment partially restores the proliferative, migratory, and reparative capacities of senescent WJ-MSCs in both *in vitro* and *in vivo* wound-healing models. Together with the observed molecular and structural modulation, these data indicate that controlled mechanical stimulation functionally augments regenerative capacity in senescent MSCs.

## 3. Conclusion

We present a microfluidic cell-reactivation platform, μ-CPR, that applies controlled hydrodynamic deformation to late-passage MSCs, eliciting a coordinated, non-genetic reactivation response. Within an optimal mechanical window, μ-CPR effectively reduced ROS levels and SA-β-Gal activity, attenuated γH2AX signals, and reinstated the expression of stemness-associated transcription factors, including OCT4, SOX2, and KLF4, concomitant with increased proliferation. These functional reactivation effects were consistently observed in WJ-MSCs and reproduced across multiple cell types beyond stem cells, demonstrating the generalizability of the approach under defined mechanical dosing conditions.

Structural and molecular analyses identified a coherent mechanistic signature. μ-CPR treatment reduced nuclear major axis length, restored actin-microtubule organization, and promoted α-actinin relocalization to focal adhesions, indicating a rebalancing of intracellular tension. Transcriptomic profiling revealed a focused reactivation signature characterized by the downregulation of pro-fibrotic ECM and SASP-linked genes, along with the enrichment of DNA repair modules. Crucially, while μ-CPR facilitates the restoration of youthful functional phenotypes, it does not fully compromise the fundamental hallmarks of stem cells, as evidenced by the preservation of intrinsic immunophenotypes and multilineage differentiation potential.

From a translational standpoint, μ-CPR offers a mechanically defined, scalable, and non-genetic approach to functionally reactivate stem cells without pharmacological or genetic intervention, making it compatible with existing cell-manufacturing workflows. Future efforts will focus on validating the durability and potency of these effects in extended culture and *in vivo* models, while refining mechanical dosing parameters to optimize the depth of the reset. Once the long-term stability and therapeutic efficacy of this mechanical stimulation are firmly established, the μ-CPR platform could pave the way toward achieving comprehensive cellular rejuvenation, ultimately transforming the landscape of regenerative medicine by decoupling functional recovery from identity alteration.

## 4. Experimental Section/Method

### Microfluidic Device Fabrication and Imaging of Cell Deformation

Microfluidic channels were designed using Autodesk AutoCAD and patterned on a 4-inch silicon wafer using conventional photolithography. Polydimethylsiloxane (Sylgard^®^ 184; Dow Corning) channel replicates were fabricated via standard soft lithography and bonded to glass slides using an O_2_ plasma treatment (CUTE; Femto Science Inc.). Cell deformation within a microchannel was visualized using a Phantom VEO710L high-speed camera (Vision Research) at frame rates of 200,000–580,000 fps and exposure times of 600 ns to 1 μs. The camera mounted on a Zeiss Axio Observer A1 inverted microscope (Carl Zeiss) captured the images.

### Cell Culture

As described in our previous studies^42^, different passages of WJ-MSCs, adipose-derived stem cells, and OOM-derived stem cells (IRB number: KUMC 2019-05-043) were cultured in MSC medium containing α-MEM (Gibco) supplemented with 10% fetal bovine serum (FBS; Peak Serum Inc.) and 1% penicillin/streptomycin (P/S; Gibco). HaCaT and BJ cells were cultured in Dulbecco’s modified Eagle’s medium-high glucose (DMEM-high) supplemented with 10% FBS and 1% P/S.

### μ-CPR Processing

Stem cell lines, HaCaT, and BJ cells were prepared at each passage and resuspended at 5 × 10^5^ cells/mL in culture medium and loaded into 10-mL syringes (BD Biosciences). PBS was loaded into a separate syringe. After connecting to the μ-CPR device, the cells were processed under controlled flow rates. Post-processing, cells were collected, centrifuged, and cultured. Cells were stabilized for 3 days prior to downstream experiments.

### Cell Proliferation Assay

Cells were plated at 5 × 10^3^ cells/well in 96-well plates (SPL Life Sciences) and maintained in MSC medium for time-course proliferation analysis. Cell proliferation was determined using the Cell Counting Kit-8 (Dojindo), following the manufacturer’s recommended protocol. Absorbance was measured at 450 nm using a Bio-RAD xMark^TM^ spectrophotometer (Bio-Rad Laboratories).

### Cell Size Measurement

WJ-MSCs from each experimental condition were cultured and photographed under a light microscope (Axiovert 40C; Zeiss). Image processing was performed using Fiji, an open-source distribution of ImageJ allowing user extensibility via Java plugins.^43^ In addition, cell size in suspension was measured using NucleoCounter^®^ NC-250™ (ChemoMetec), according to the manufacturer’s protocol.

### ROS Generation Assessment using H_2_DCF-DA

To determine intracellular ROS accumulation, WJ-MSCs were cultured in an MSC medium and grown to confluence. Cells were washed and incubated with 10 μM 2’,7’-dichlorodihydrofluorescein diacetate (H_2_DCF-DA, Invitrogen) reagent for 30 min, as described by the manufacturer. After incubation, the solution was replaced with PBS, and the WJ-MSCs were directly visualized using a fluorescence microscope (Nikon). In addition, the fluorescence intensity of the H_2_DCF-DA probe was imaged using a fluorescence microscope, and the mean fluorescence intensity was analyzed using ImageJ.

### Reverse Transcription-Polymerase Chain Reaction

Total RNA was isolated from each group of WJ-MSCs using Labozol Reagent (LaboPass; Cosmo Genetech), as defined by the manufacturer. RNA concentration and purity were assessed using a NanoDrop spectrophotometer (NanoPhotometer^®^ N50, IMPLEN). cDNA synthesis was performed using 2 μg total RNA using the M-MuLV reverse transcription kit (Labopass) and oligo (dT) primers. HiPi Real-Time PCR 2× Master Mix (Elpis Biotech) was used to measure the relative gene expression levels between the control and samples treated with μ-CPR. The primers used in this study are listed in **Table S1**. The methods were performed following the manufacturer’s protocol. The relative expression results were normalized to GAPDH (internal control), and the fold change (FC) in gene expression was calculated using the comparative cycle time method.

### Immunofluorescence Staining

WJ-MSCs were fixed with 4% paraformaldehyde (PFA), permeabilized with PBST [0.2% (vol/vol) Triton X-100 in PBS], and blocked with 10% (vol/vol) bovine serum albumin in PBST. Specimens were incubated overnight at 4 °C in PBST with 1% bovine serum albumin and the following primary antibodies: OCT4 (sc-9081; Santa Cruz Biotechnology), SOX2 (sc-20088; Santa Cruz Biotechnology), SSEA4 (sc-21704; Santa Cruz Biotechnology), phospho-histone H2A.X (2577S; Cell Signaling Technology), actin (sc-8432; Santa Cruz Biotechnology), α-actinin (sc-17829; Santa Cruz Biotechnology), and α-tubulin (sc-23948; Santa Cruz Biotechnology). The WJ-MSCs were washed three times with PBST and incubated with secondary antibodies (Alexa Fluor goat anti-mouse 488, A-11001, 1:500 dilution, or Alexa Fluor goat anti-rabbit 546, A-11035, 1:500 dilution; Thermo Fisher Scientific) for 2 h at room temperature (RT), followed by three additional washes with PBST. For staining of anti-histone H3 (tri-methyl K27, Millipore), cells were incubated with Alexa Fluor 488 (Invitrogen) goat anti-rabbit IgG (H + L), cross-adsorbed secondary antibody. Samples were imaged using a fluorescence microscope (LSM900 with Airyscan 2; Zeiss).

### Western Blotting

Cell lysates were prepared in extraction buffer composed of 100 mM Tris-HCl (pH 7.5), 1% Triton X-100 (Sigma-Aldrich), 10 mM NaCl, 10% glycerol (Amresco), 50 mM sodium fluoride (Sigma-Aldrich), 1 mM phenylmethylsulfonyl fluoride (PMSF; Sigma-Aldrich), 1 mM p-nitrophenyl phosphate (Sigma-Aldrich), and 1 mM sodium orthovanadate (Sigma-Aldrich). The lysates were centrifuged at 16,000 g for 15 min at 4 °C, and the supernatant was carefully transferred into a new E-tube for protein quantification using a BCA protein assay kit (Thermo Fisher). The proteins (15 μg) were separated using 8–12% sodium dodecyl sulfate-polyacrylamide gel electrophoresis and then transferred onto nitrocellulose membranes (Bio-Rad). Membranes were blocked using 5% skimmed milk in Tris-buffered saline for 1 h, then incubated overnight at 4 °C with the appropriate primary antibodies (Abs): OCT4 (Santa Cruz Biotec, sc-5279), SOX2 (CST, 23064), SSEA4 (Santa Cruz Biotec, sc-21704), PCNA (CST, 2586), and anti-β-actin (CST, 4970S). Subsequently, the membranes were incubated with secondary antibodies (anti-mouse or anti-rabbit IgGs) conjugated with horseradish peroxidase (Santa Cruz Biotechnology) for 1 h at RT. Protein signals were visualized using an enhanced chemiluminescence kit (Amersham Biosciences) and a ChemiDoc^TM^ Imaging System (Bio-RAD, 17001401).

### SA-β-Gal Assay

The P6, P11, and P18 μ-CPR cells were prepared, and the SA-β-Gal assay was performed as previously described.^44^ Cells were cultured in a 6-well cell culture plate (SPL Life Sciences). Upon reaching 80% confluence, they were washed with PBS, fixed for 15 min with 2% PFA and 0.2% glutaraldehyde. Subsequently, the cells were incubated for 15 h at 37 °C (non-CO_2_ regulated) with freshly prepared SA-β-Gal staining solution containing 200 mM citric acid/sodium phosphate, 100 mM K_4_[Fe (CN)_6_] • 3 H_2_O, 100 mM K_3_[Fe (CN)_6_], 5 M NaCl, 1 M MgCl_2_, and 50 mg/mL X-Gal. Blue-stained SA-β-Gal-positive cells a were observed under a light microscope.

### RNA-seq analysis

RNA samples from WJ-MSCs cultured under each condition were isolated from the cell pellets using TRIzol^TM^ Reagent (Invitrogen). The RNA extracts were sent to Ebiogen Inc. for library preparation and RNA sequencing. The libraries were prepared using the NEBNext Ultra II Directional RNA-Seq Kit (New England Biolabs, Inc.) and sequenced on the Illumina NextSeq 500/550. Raw RNA data were converted to Excel-based DEG Analysis (ExDEGA) (Ebiogen Inc.) files for efficient analysis. Specifically, the adapter sequences and low-quality reads (quality score < 20) were removed using FASTX_Trimmer (http://hannonlab.cshl.edu/fastx_toolkit/). The trimmed sequencing file was mapped to the human reference genome hg19 using TopHat2.^45^ The gene expression levels were estimated using Cufflinks,^46^ followed by differential expression analysis using EdgeR.^47^ DEGs from the μ-CPR group were classified relative to the control. DEGs were defined using thresholds of *P* ≤ 0.05, |log FC| ≥ 1. PCA was performed on the normalized data (z-score) transcriptional expression data using XLSTAT software, version 2021.4 (Addinsoft, 2021).

### Heatmap Analysis

RNA-seq data from the μ-CPR-exposed cells (P18) were analyzed against the corresponding control groups, referencing the age-related gene list from the GenAge and CellAge databases (HAGR, Human Aging Genomic Resources, https://genomics.senescence.info/genes/).^48^ Gene sets from each group were identified and categorized using KEGG and REACTOME databases. Then, classified genes were ordered based on hierarchical clustering using Euclidean distance and the average linkage method in MultiExperiment Viewer (MeV) v4.9.0, The TM4 Development Group, http://www.tm4.org) and Prism (version 7 GraphPad Software)^49^ to generate heatmaps.

### Antibody Array Assay

The Antibody Array slide (RayBiotech, Norcross, GA) was dried for 2 h at RT and incubated with 400 µL of blocking solution for 30 min. After decanting the blocking buffer from each sub-array, 400 µL of diluted samples were added and incubated for 2 h at RT. Following this, the samples were decanted, and each array was washed thrice with 800 µL of 1X wash buffer I for 5 min with gentle shaking. The glass chip assembly was placed into the container, and submerged in sufficient 1X wash buffer for 10 min with shaking performed twice. An additional washing step was carried out using 1X wash buffer. 1X biotin-conjugated anti-cytokine antibodies were prepared and incubated for 2 h at RT with gentle shaking, followed by washing with 150 µL of 1X wash buffer at RT with shaking. Then, 1X Cy3-conjugated streptavidin stock solution was added and incubated for another 2 h at RT with gentle shaking, followed by two washes with 1X wash buffer I for 10 min each. After washing, the slide was rinsed with deionized water using a plastic wash bottle and centrifuged at 1,000 rpm for 3 min to remove excess water. The slide was scanned with a GenePix 4100A Scanner (Axon Instruments) and allowed to dry completely within 24–48 h. Scanning was conducted at 10 µm resolution with optimized laser power and PMT settings. The scanned images were grid-divided and quantified using GenePix Software (Axon Instruments). Subsequently, protein data were annotated using the UniProt database, and data mining and visualization were performed using ExDEGA (Ebiogen Inc.).

### *In vitro* Cell Scratch Assay

To assess migration, WJ-MSCs were seeded in 6-well plates (SPL Life Sciences) and cultured in MSC medium until confluence. Cells were treated with 10 μg/mL mitomycin C (Sigma-Aldrich) for 2 h at 37 °C to inhibit proliferation, washed, and scratched with a 1,000-μL pipette tip in serum-free MSC medium. Wound closure was monitored over time (biological triplicates) and quantified using TScratch software.^50^

### In Vivo Wound Healing Assay

To analyze the *in vivo* wound healing capacity, we used six-week-old BALB/c nude female mice (CAnN.Cg-Foxn1nu/CrljOri SPF/VAF Immunodeficient mice, ORIENT BIO Animal Center). The *in vivo* experiments were performed following approval from the Institutional Animal Care and Use Committee (IACUC) at Konkuk University (approval no.: KU20132). One week before the experiment, mice were acclimated in a well-ventilated room with controlled temperature and humidity, under a 12 h light/12 h dark cycle, and had *ad libitum* access to food and water. They were divided into six groups as follows: (1) control wound group, (2) PBS-treated group, (3) P6-treated group, (4) P11-treated group, (5) P18-treated group, and (6) P18 μ-CPR-treated group (n = 4 mice per group). Before creating the *in vivo* wound, all mice were anesthetized via intraperitoneal injection of 60 mg/kg of alfaxalone-xylazine (Alfaxan; Careside, Rompun; Elanco). Mice were anesthetized, and wounding was performed using a sterile biopsy punch (diameter of 8 mm; Kai Industries) to create two full-thickness skin wounds on the back of each animal. Mice were then injected subcutaneously at four points around each wound with either PBS (for the control group) or 2 × 10^6^ cells/mL of WJ-MSCs suspended in 100 μL of PBS. To protect against contamination, wounds were sealed with silicone (0.5 mm T), Tegaderm tape (1622 W), and a bandage (DUPOL). The progress of wound closure was monitored using a digital camera. The size of the wounds was measured using a 30-cm ruler and recorded using a camera in a time-dependent manner. Additionally, scar formation was assessed two weeks post-wounding. On days 6 or 12 after cell injection, the mice were sacrificed, and the skin tissues around the wounds were excised, fixed in 4% PFA, and dehydrated using increasing alcohol concentrations, followed by paraffin embedding. Subsequently, the tissues were cut perpendicular to the wound surface into 4-µm-thick tissue sections and mounted on slides pre-coated with 0.1% w/v poly L lysine (Sigma-Aldrich). The sections were subjected to hematoxylin and eosin staining to visualize tissue lesions and regeneration and evaluate the degree of re-epithelialization. Masson’s trichrome staining was performed to estimate the rate of collagen synthesis, while immunohistochemistry staining was performed with PCNA (CST, 2586) and β-catenin (Santa Cruz Biotechnology, sc-7963) to evaluate the regeneration effect. Digital slide scanning was conducted using a 3DHISTECH scanner.

### Statistical Analysis

Statistical analysis was performed using GraphPad Prism (GraphPad Software, version 7). Experiments were performed in biological triplicate unless noted otherwise. Statistical significance was assessed using Student’s t-test or two-way ANOVA, as appropriate. In all figures, *P*-values are marked with asterisks (*P* < 0.05 (*), *P* < 0.01 (**), *P* < 0.001 (***), *P* < 0.0001 (****)).

## Acknowledgements

This research was supported by the Samsung Research Funding & Incubation Center of Future Technology (SRFCIT1802-03 to A.J.C. and S-G.C.), the Korea Fund for Regenerative Medicine (KFRM) grant funded by the Korean government (Ministry of Science and ICT and Ministry of Health and Welfare; 24A0203L1 and 25B0101L1 to S-G.C.), and National Research Foundation of Korea (NRF) grants funded by the Korean government (MSIT; Ministry of Science and ICT; RS-2023-00218543 to A.J.C.).

## Data Availability Statement

Data from this study are available from the corresponding authors upon reasonable request.

## ToC

### Microfluidic Mechanical Reactivation of Aged Stem Cells

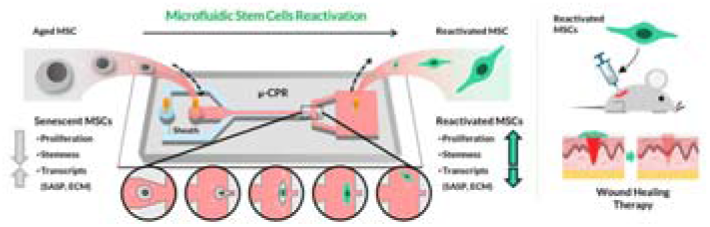

This study introduces μ-CPR, a microfluidic technology that revitalizes aged stem cells using controlled mechanical pressure instead of chemicals. By improving the cell’s internal structure and function, μ-CPR restores youthful growth and healing abilities while keeping the cell’s original identity intact. This safe, non-genetic method provides an effective strategy for producing high-quality stem cell therapies to treat aging-related conditions.

