## Supplementary material for "Microfluidic Mechanical Reactivation of Aged Stem Cells": SI

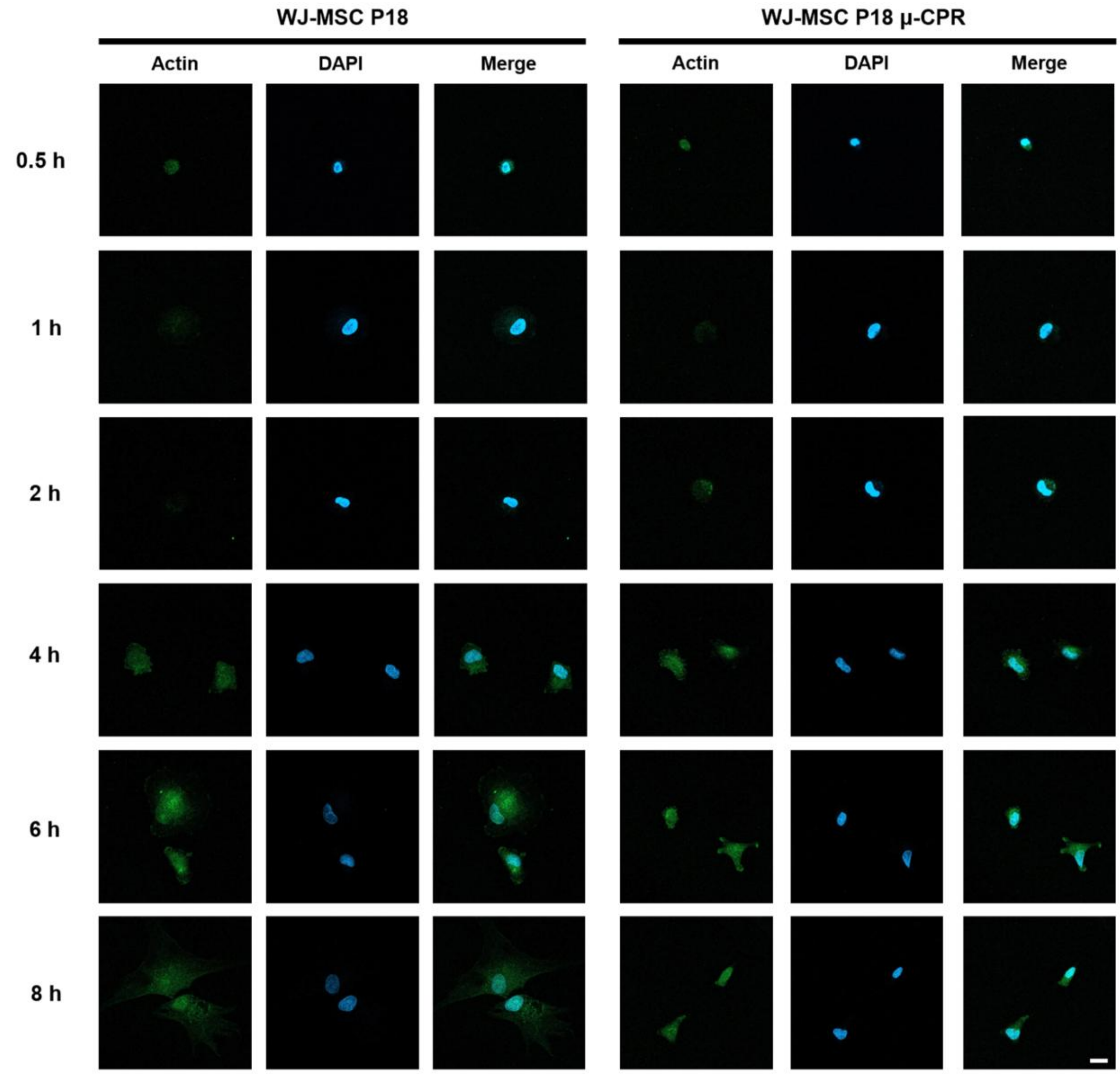


**Figure S1.** Time-dependent comparative analysis of actin expression in late-passage WJ-MSCs (P18) and μ-CPR-treated WJ-MSCs (P18 μ-CPR) via immunofluorescence. Scale bar = 20 μm.

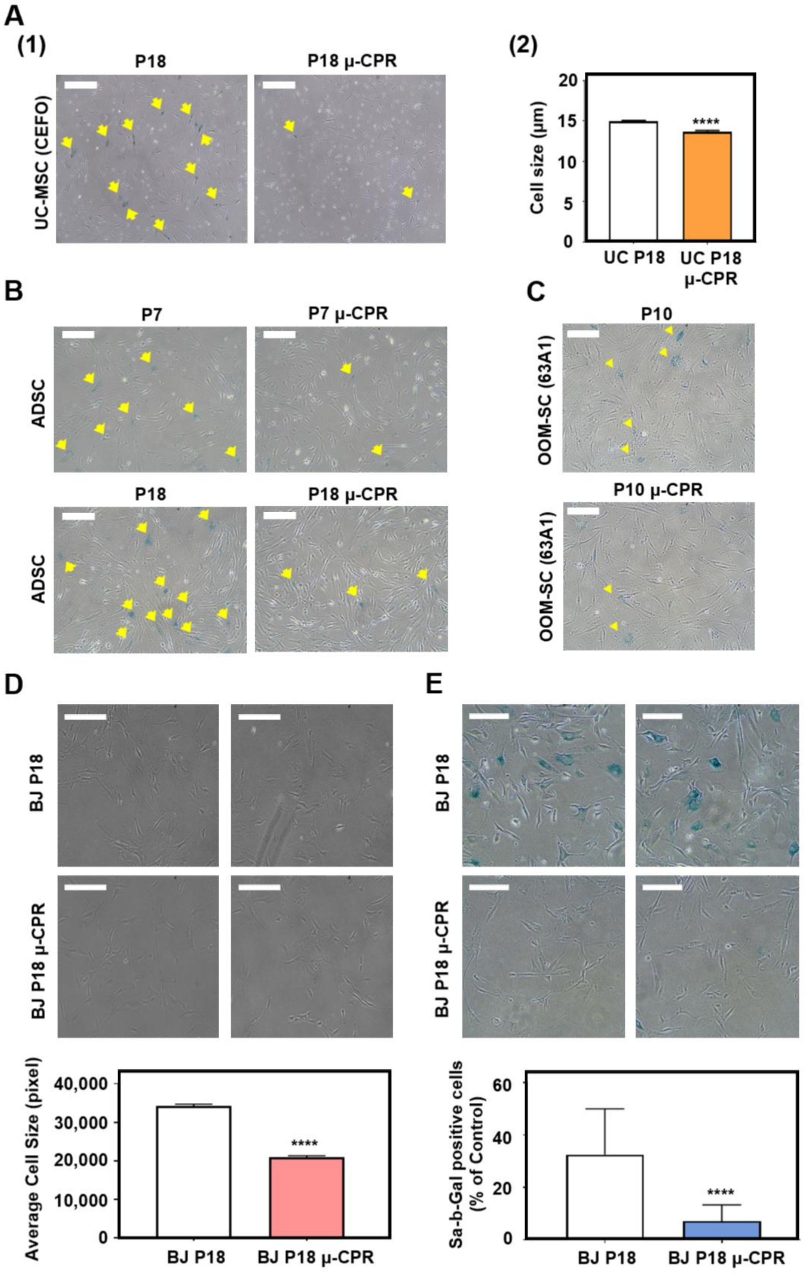


**Figure S2. Comparative analysis of senescence markers following μ-CPR treatment in various cell types**. **(A)** Umbilical cord-derived mesenchymal stem cells (UC-MSCs) and μ-CPR-treated UC-MSCs were analyzed for senescence-associated β-galactosidase (SA-β-Gal) staining and cell size. **(B)** Senescence-positive cells in adipose-derived stem cells (ADSCs) at passages 7 and 18 were assessed via SA-β-Gal staining. Scale bar = 500 μm. **(C)** Comparative analysis of SA-β-Gal-positive cells in orbicularis oculi muscle (OOM) tissues of different ages. Scale bar = 500 μm**. (D)** Late-passage human fibroblast (BJ) cells (P18) and μ-CPR-treated BJ cells (P18 μ-CPR) were compared for cell size (measured using ImageJ) and SA-β-Gal positivity. Scale bar = 500 μm. Graphs show the SA-β-Gal-positive cells. Data represent the mean ± SEM of three independent experiments. Statistical analysis was performed using one-way ANOVA; **P* < 0.05, ***P* < 0.01, ****P* < 0.001, *****P* < 0.0001.

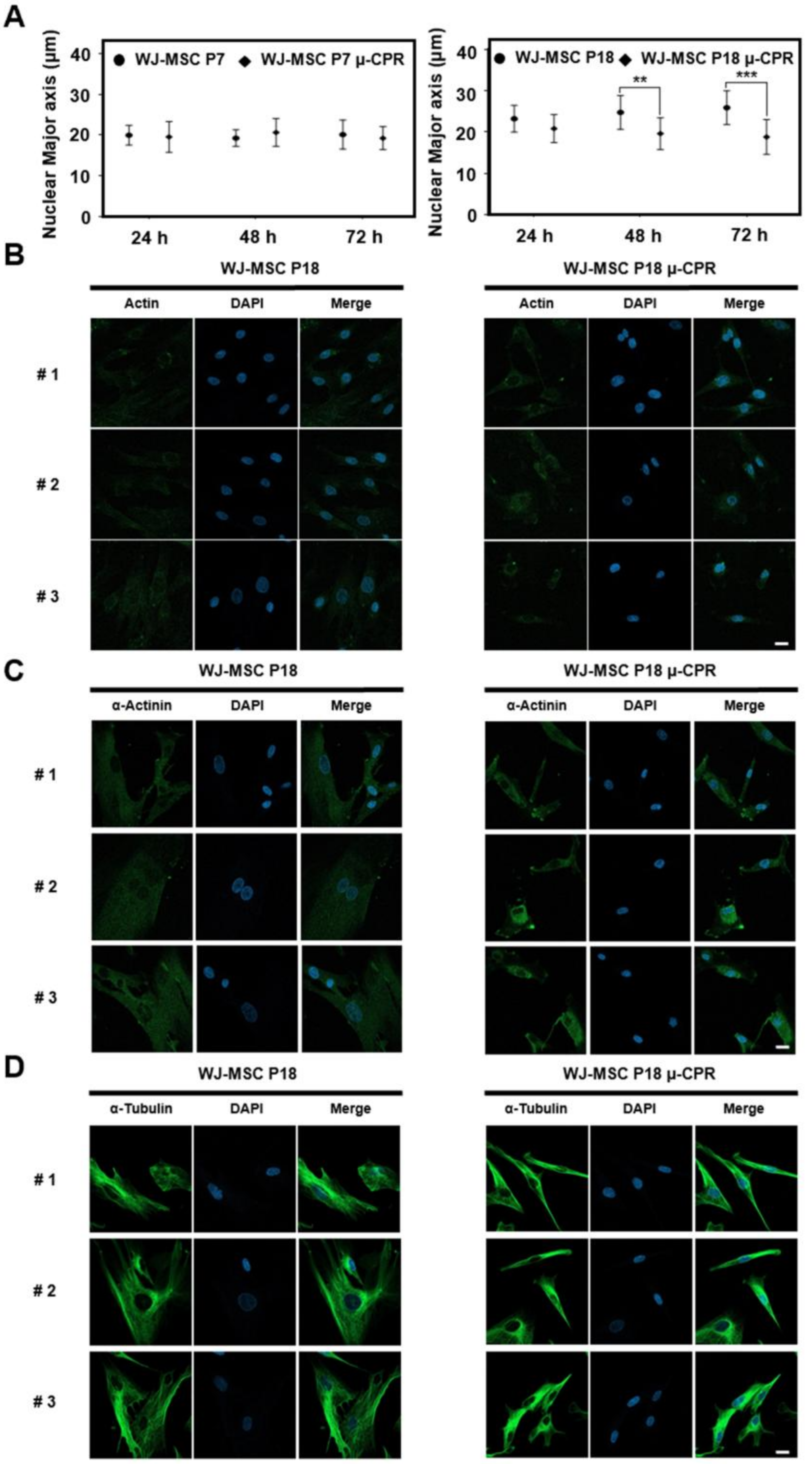


**Figure S3. Comparison of cytoskeletal organization in late-passage WJ-MSCs (P18) with and without μ-CPR treatment. (A)** Quantification of nuclear size showing the effects of μ-CPR and senescence. Left: comparison between WJ-MSCs P7 and P7 μ-CPR; Right: nuclear size changes in WJ-MSCs P18 following μ-CPR. **(B–D)** Immunocytochemical analysis of cytoskeletal components in WJ-MSCs P18 and P18 μ-CPR: **(B)** actin, **(C)** α-actinin, and **(D)** α-tubulin. Scale bar = 20 μm.

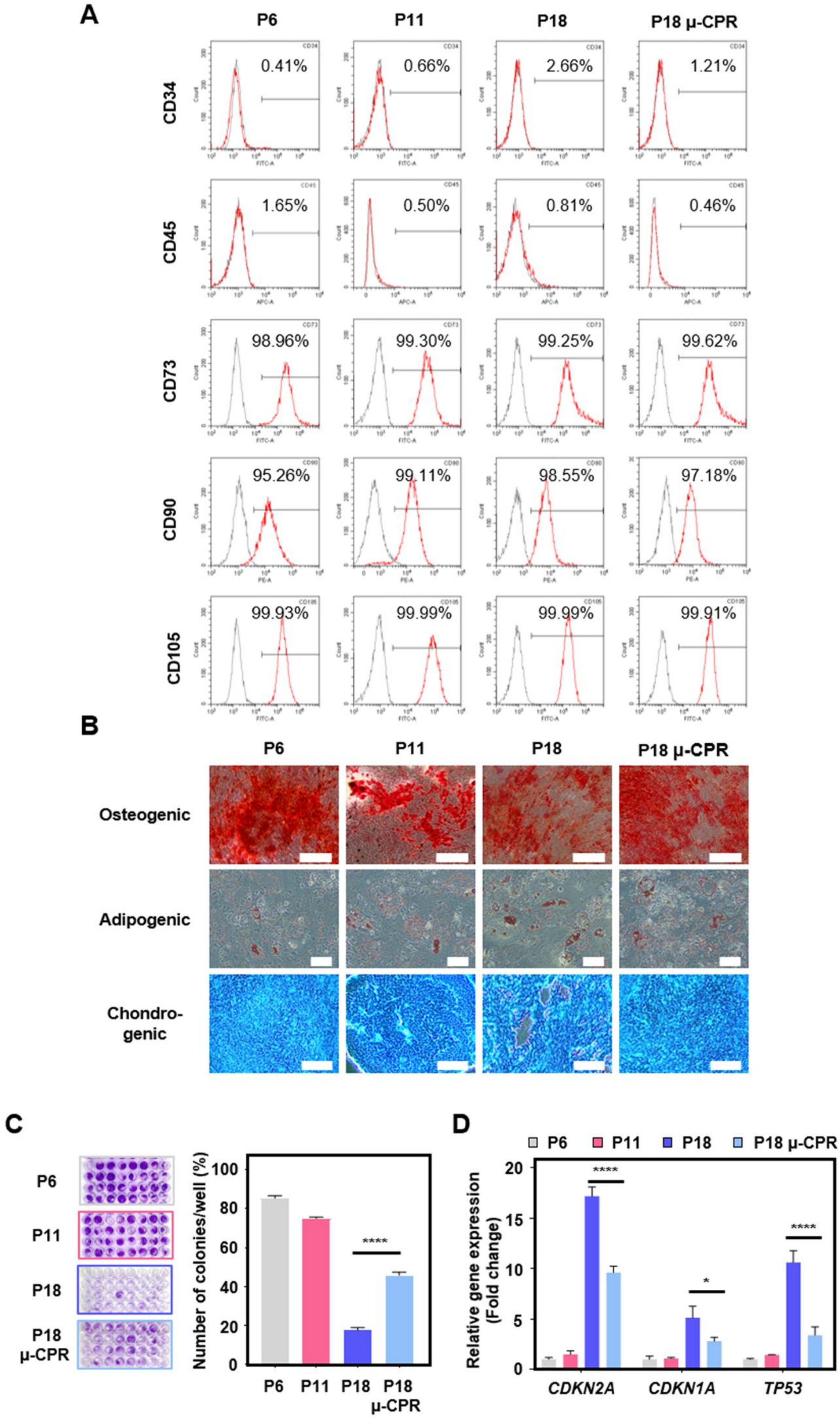


**Figure S4. Evaluation of the protective effects of μ-CPR against replicative senescence in long-term cultured WJ-MSCs. (A)** Comparative analysis of cell surface marker expression across P6, P11, P18, and P18 μ-CPR cells. **(B)** Representative images of trilineage differentiation (osteogenic, adipogenic, and chondrogenic) in WJ-MSCs at P6, P11, P18, and

P18 μ-CPR (scale bar = 500 μm). **(C)** Comparison of colony-forming unit (CFU) formation among P6, P11, P18, and P18 μ-CPR cells. **(D)** Quantitative analysis of senescence-related genes (*CDKN2A* (p16), *CDKN1A* (p21), and *TP53*) in P6, P11, P18, and P18 μ-CPR cells. Data represent the mean ± SEM of three independent experiments. Statistical analysis was performed using one-way ANOVA; *$P < 0.05$, **$P < 0.01$, ***$P < 0.001$, ****$P < 0.0001$.

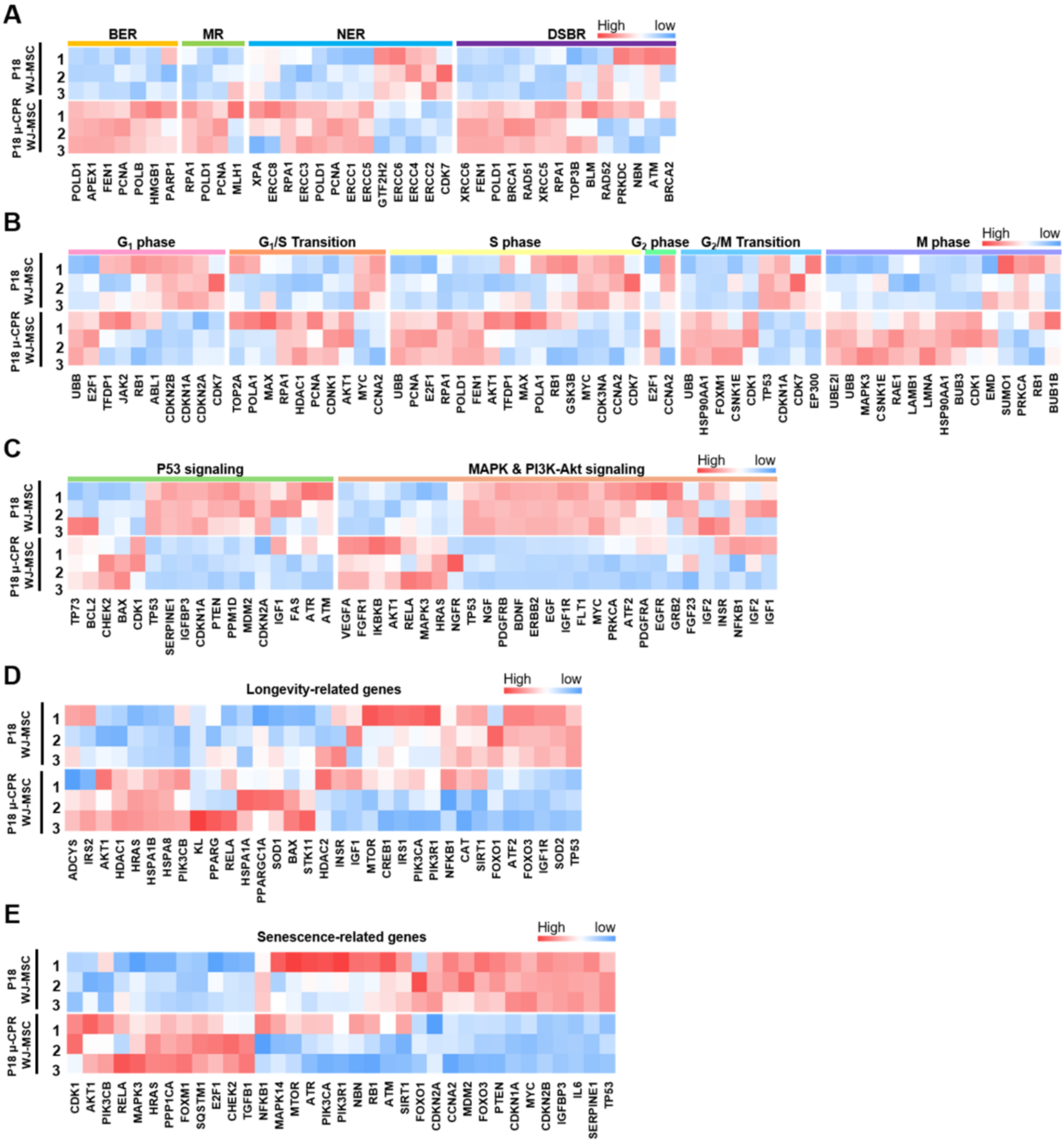


**Figure S5. Bioinformatic analysis of senescent WJ-MSCs following µ-CPR treatment**. Hierarchical clustering heatmaps compare gene expression patterns between P18 WJ-MSCs and P18 µ-CPR across multiple pathways. **(A)** DNA damage response (DDR), **(B)** cell cycle, **(C)** p53 and MAPK/PI3K-Akt signaling pathways, **(D)** longevity-associated genes, and **(E)** senescence-associated genes. Heatmaps were generated using Euclidean distance clustering with MeV software. Cell colors indicate changes in gene expression based on z-score thresholds.

**Table S1**. Primer sequences for stemness-related genes used in reverse transcription-quantitative polymerase chain reaction (RT-qPCR) experiments

| **Gene** | **Primer** | **Product Size (bp)** | **ACCESSION No.** |
|---|---|---|---|
| ***GAPDH*** | F-5'-GTC TCC TCT GAC TTC AAC AGC G-3'<br>R-5'-ACC ACC CTG TTG CTG TAG CCA A-3' | 131 | NM_001357943.2 |
| ***OCT4*** | F-5'-CCT GAA GCA GAA GAG GAT CAC C-3'<br>R-5'-AAA GCG GCA GAT GGT CGT TTG G-3' | 106 | NM_001173531.3 |
| ***SOX2*** | F-5'-GCT ACA GCA TGA TGC AGG ACC A-3'<br>R-5'-TCT GCG AGC TGG TCA TGG AGT T-3' | 135 | NM_003106.4 |
| ***KLF4*** | F-5'-CAT CTC AAG GCA CAC CTG CGA A-3'<br>R-5'-TCG GTC GCA TTT TTG GCA CTG G-3' | 156 | NM_001314052.2 |
